# Fold or flop: quality assessment of AlphaFold2 predictions on whole proteomes

**DOI:** 10.64898/2025.12.19.695427

**Authors:** Edoardo Sarti, Frédéric Cazals

## Abstract

**Motivation:** Reliability of `AlphaFold2` predictions is mainly assessed using the predicted Local Distance Difference Test (pLDDT). For model organisms, 30–40% of residues fall into the low-confidence pLDDT range. Moreover, pLDDT sometimes fails to flag physically implausible structures. This raises two questions: can more robust reliability indicators be identified, and do unreliable predictions share common structural or biophysical features?

**Results:** We characterize protein structures through histograms of per-residue neighbor counts, and use the Wasserstein principal component analysis to define the arity map, and lightweight and informative 2D embedding of proteins in a dataset. Using `AlphaFold-DB`, we show that the arity map reveals three structurally and biophysically distinct populations (well-folded proteins, intrinsically disordered proteins, and physically implausible predictions). We also use our packing based encoding at the residue level to define `abstraqt` (Arity-Based STRuctural Arrangement Quality assessmenT), a per-residue scoring function complementing the pLDDT, assigning low scores to hallucinated helices and distorted beta strands while correctly scoring native like predictions.

**Availability:** The code to compute arity maps is available within the [Structural Bioinformatics Library](https://sbl.inria.fr/). See: [AlphaFold analysis](https://sbl.inria.fr/doc/Alphafold_analysis-user-manual.html), and also [Documentation](https://sbl.inria.fr/doc/), [Applications](https://sbl.inria.fr/applications/), [Installation guide](https://sbl.inria.fr/doc/sbl-installation-guide.html).

The code and data for rerunning analyses are made available from [AlphaFold analysis](https://sbl.inria.fr/doc/Alphafold_analysis-user-manual.html), and [Data](https://sbl.inria.fr/data/AlphaFold-assessment).

## 1 Introduction

`AlphaFold2` has revolutionized protein structure prediction, which in favorable cases achieves high accuracy comparable to experimental methods [1]. Its companion resource, the AlphaFold Protein Structure Database (`AlphaFold-DB`) now hosts more than 200 million predicted structures generated by `AlphaFold2`. `AlphaFold-DB` has rapidly become the primary reference for structural information on `UniProtKB` sequences, serving both as a comprehensive repository and as a springboard for derivative resources [2, 3] extending even more the range of predicted structures.

Each amino acid of any `AlphaFold2` prediction is assigned a confidence score known as pLDDT (predicted Local Distance Difference Test). It is derived from the Local Distance Difference Test (LDDT [4]), a superposition-free score that evaluates how well interatomic distances in a predicted structure match those in a reference. During training, `AlphaFold2` was supervised using known X-ray and CryoEM experimental structures, with a dedicated confidence head in the network predicting the expected LDDT score for each residue. The resulting pLDDT values are normalized from 0 to 1, where higher values indicate greater confidence in local geometry. For simplicity, four confidence categories have been defined based on pLDDT values: very high (0.9 ≤ pLDDT, dark blue), high (0.7 ≤ pLDDT *<* 0.9, cyan), low (0.5 ≤ pLDDT *<* 0.7, yellow), and very low (pLDDT *<* 0.5, orange). Importantly, pLDDT reflects local accuracy only, without assessing global domain orientations or inter-chain relationships, which are instead captured by complementary metrics such as Predicted Alignment Error (`PAE`).

The structural features within `AlphaFold2` predictions have been examined using the `Geometricus` algorithm [5], which represents protein structures through a series of steps: (i) a decomposition of the protein into structural fragments (based on *k*-mers or spherical neighborhoods), (ii) a discrete encoding of invariants (moments) of these fragments into shape-mers, and (iii) a counting step of shape-mers yielding an embedding of the protein. By clustering these embeddings – using non-negative matrix factorization – across 21 predicted proteomes from `AlphaFold-DB`, 250 structural clusters were identified, including 20 major clusters or superfamilies [6].

The pLDDT metric has been thoroughly analyzed, revealing its effectiveness as an indicator of result robustness. It has been observed that the low-pLDDT regions in `AlphaFold2` predictions correlate with observed or predicted intrinsically disordered regions (IDRs) [7]. Indeed `AlphaFold2` outperforms state-of-the-art methods such as `IUPred2` on disordered region detection on a benchmark of `DisProt` targets and PDB controls [6]. Also, high mean pLDDT values are associated with minimal structural differences between `AlphaFold2` and `trRosetta` models [6].

For the 49 model organisms whose entire proteome have been predicted and deposited on `AlphaFold-DB`, 30% to 40% of all amino acids fall within the low or very low pLDDT confidence categories (Table S1; Figures S1 to S3.). Although a considerable fraction could be explained by intrinsically disordered regions (IDRs), many low pLDDT amino acids are situated in the bulk, or are not part of low pLDDT segments. Moreover, pLDDT does not correlate with the local flexibility of globular proteins [8]. Even in regions predicted with high confidence, `AlphaFold2` models can disagree with experimentally determined electron density maps concerning domain orientations, local backbone, and side-chain conformations [9]. Finally, `AlphaFold2` lacks robustness to minor, biologically insignificant sequence perturbations. Adversarial sequences with only five altered residues can lead to drastically different 3D structures, yet `AlphaFold2` often predicts these with similar confidence to original sequences [10].

Despite the evidence of faults in a small yet significant fraction of `AlphaFold2` predictions, reports have so far remained anecdotal and linked to the context of specific studies. Is there a common denominator between `AlphaFold2` incorrect predictions? Could we use this information to create a better confidence measure?

### Contributions

We make three contributions. First, we characterize protein structures through histograms of per-residue neighbor counts, and use the Wasserstein principal component analysis (based on the Wasserstein distance 𝒲_2_) to define and lightweight and informative 2D embedding of proteins in a dataset. Second, we show that the arity map reveals three structurally and biophysically distinct populations (well-folded proteins, intrinsically disordered proteins, and physically implausible predictions) and exposes limitations of existing quality metrics that fail to discriminate between these groups. Third, we extend the same neighbor count histogram framework to the residue level and introduce `abstraqt` (Arity-Based STRuctural Arrangement Quality assessmenT), a per-residue scoring function trained on a curated dataset of plausible and implausible structural contexts. `abstraqt` complements pLDDT by assigning low scores to hallucinated helices and distorted beta strands while correctly scoring both globular and non-globular well-determined structures.

## 2 Methods

### 2.1 Packing analysis

#### 2.1.1 Global packing analysis: arity and arity signature

A natural parameter to identify compact regions in proteins in the number of neighbors of a *C*_*α*_ carbon. Let the *arity* of a *C*_*α*_ carbon be the number of neighboring *C*_*α*_s within a distance range *r*. As classically done, we take *r* = 10Å to account for non-covalent contacts–Sec. S1.0.1. We note in passing that such neighbors are sufficient to robustly infer protein domains using a direct application of spectral clustering [11]. Assuming the polypeptide chain has *n* amino acids, define – Fig. S1:

- *L*_*a*_ = {*a*_*n*_, …, *a*_*n*_}: arities of the *n* individual *C*_*α*_ carbons;
- *L* = [*A*_1_, …, *A*_*m*_]: unique arities sorted by increasing value;

The histogram of arities is denoted 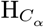.

Our goal is to analyze a set of structures associated with polypeptide chains {*P*_*i*_*}*, say from `AlphaFold-DB`. We therefore define:

##### Definition. 1

*(Arity signature Sig*_*i*_(*P*) *of a polypeptide chain P*_*i*_.*) The discrete probability measure associated with the normalized arity histogram is denoted ν*_*i*_, *and the associated cumulative distribution function F*_*i*_. *We also define the associated quantile function*

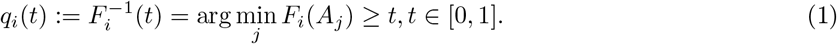

*Consider a list P*_*K*_ = {*p*_1_, …, *p*_*K*_} *of increasing quantiles/percentiles. The C*_*α*_ signature *is the increasing sequence Sig*_*i*_(*P*) = {*q*_*i*_(*p*_1_), …, *q*_*i*_(*p*_*K*_)}.

The arities can also be used to *build* the protein incrementally, by inserting the a.a. by increasing arity. Upon inserting one amino acid, we connect it to its neighbors along the sequence if already present, a classical task carried out with the Union-Find algorithm [12]. As we will see, the evolution of the number of connected components is a signature of the overall packing.

An obvious instantiation of the previous framework consists of using two quantiles/percentiles *p*_1_ and *p*_2_ to define a 2D embedding of the structures of interest, that is:

##### Definition. 2

*Given two quantiles/percentiles p*_1_ *< p*_2_, *the* arity map *is the map whose x and y axis are the arities at p*_1_ *and p*_2_. *A* bin/cell *of the map hosts all structures with a prescribed arity signature*.

We now develop the proper mathematical apparatus to justify this definition and the 2D embedding defined from two quantiles.

##### Remark 1

*Populated bins in the arity map are naturally above the diagonal y* = *x. To compare two arity maps, the lower region will be used, with mirrored coordinates*.

#### 2.1.2 Wasserstein PCA on arity signatures

The arity histogram or the associated discrete measure *ν*_*i*_ encodes global packing properties, and we can compare two such distributions using the p-th Wasserstein distance 𝒲_*p*_. For *p* ≥ 1, this distance boils down to comparing the quantile functions, a consequence of the usual uncrossing argument [13, Thm 2.18] or [14]:

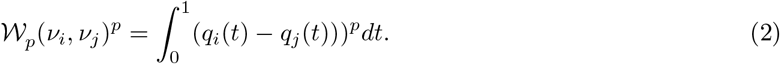

In the sequel, we consider *p* = 2 which enables using the framework of functional data analysis in the Wasserstein space, in which case the Wasserstein center of mass of *n* quantile functions satisfies [15, 16]:

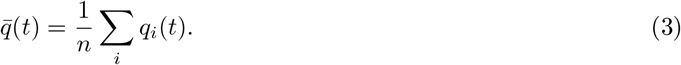

Since the quantile functions can be centered and we can compute the dot product of the centered versions, we can directly perform PCA, known as *Wasserstein PCA*.

Practically, assume *q*_*i*_ is discretized as a 1 × *T* vector of quantile values, with *T* the number of quantiles chosen. (In practice, we use percentiles so that *T* = 100.) The data matrix *Q* for *n* proteins is the *n* × *T* matrix obtained by stacking these vectors. Let **1**_**n**_ be the *n* × 1 vector of ones. The mean quantile function (Eq. 3) reads as

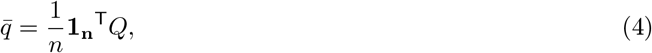

whence the data centered matrix

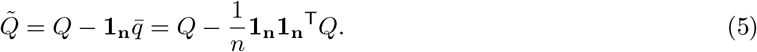

We then compute the following covariance matrix which is used to perform PCA

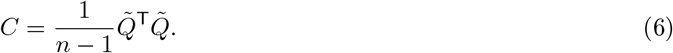

As usual, a low dimensional embedding of the data points (the arity distributions here) can be obtained using the eigenvectors associated with the leading eigenvalues of the covariance matrix. The reader is referred to Sec. 3.1 for the justification of Def. 2.

#### 2.1.3 Local packing analysis

In order to characterize a single residue in an analogue fashion as what we described for complete structures, we calculate its *local arity histogram*, which contains the normalized neighbor counts for each of its neighboring residues. Consistently with the previous section, we define the distance between two of such histograms as the Wasserstein distance 𝒲_2_.

### 2.2 Backbone geometric analysis

We use a classical geometrical approach to detect unusual backbone conformations of secondary structure elements (SSEs). More precisely, to define the SSE center line with minimal complexity, we first calculate the center of mass of three successive intervals of *N C*_*α*_ shifted by *k* residues, where *N* = 3, *k* = 2 for beta sheets and *N* = 4, *k* = 4 for alpha helices. We then plot a B-spline *C*(*t*) of degree 3 through the said center of masses.

The local curvature radius at any given point is then calculated with the usual formula [17]:

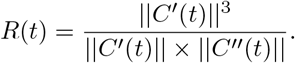

Using the residue-wise curvature value as decision parameter, we then define a secondary structure element as pathological according to the following criteria:

- For alpha helices, whether the curvature radius of the helix center line over 12 residues is lower than 30 Å;
- For beta strands, whether the curvature radius of the beta sheet center line over 4 residues is lower than 5 Å, or whether the torsion of the beta sheet over 4 residues is greater than 40 degrees.

### 2.3 abstraqt: an arity-based structure quality assessment method

#### 2.3.1 Method: automatic detection of implausible structural motifs

Within the `AlphaFold-DB` predictions with non-standard arity values, we find an over representation of certain unlikely or nonphysical structural motifs. The two predominant ones are 100-to 300-residue long helices (often containing narrow bends), and beta solenoid structures. These structures have been observed and catalogued as two recurrent `AlphaFold2` hallucinations [18], which are frequently assigned with pLDDT *>* 50 (High or Very High confidence).

Leveraging the framework presented in the previous section, we describe `abstraqt` (Arity-Based STRuctural Arrangement Quality assessmenT), a simple, residue-wise scoring function that complements the pLDDT measure detecting and down scoring implausible structural motifs. `abstraqt` is a kNN classifier [19, 20] trained over a number of positive (well-folded or intrinsically disordered residues) and negative cases (misfolded or incorrectly-predicted disordered residues): for each binary-labeled residue of the training set, the *local arity histogram* is calculated (Section 2.1.3). Then, a Ball tree [21] is constructed using the 𝒲_2_ distance to perform efficient nearest neighbor search.

For each residue of a target protein structure, `abstraqt` calculates its local arity histogram, queries the Ball tree and averages the binary labels of the *k*(= 100) nearest training set histograms.

#### 2.3.2 Training set

In order to minimize biases in the kNN classifier, we build an equilibrated dataset representing correctly- and incorrectly-predicted ordered and disordered regions. Our 100,000 amino acid training set is thus divided into five balanced parts of 20,000 amino acids each, each receiving the label 1 if the residue is found in a physically plausible context and 0 otherwise:

- Residues belonging to `CATH` [22] domains within the context of the protein
- Residues belonging to `CATH` domains where the domain was extracted from the protein
- Residues forming implausible secondary structures.
- Residues forming disordered segments of at least 10 amino acids that were experimentally confirmed in `DisProt` [23].
- Residues forming disordered segments of at least 10 amino acids that were not confirmed by the disorder predictor `AIUPred` [24].

##### Positive-ordered: experimentally-determined folded regions from CATH

In order to sample uniformly from experimentally determined folds, we take `CATH` [22] as our reference classification and sample from its 9,798 non redundant domains with less than 20% sequence similarity. For each residue extracted from `CATH` domains, arity is calculated in two ways: keeping into account the whole reference structure where the domain has been identified, and keeping into account only the residues within the domain. This double annotation helps training recognition of typical structural motifs even when the structure is not complete, as in the case of predicting the structure of a single chain of a multimer.

##### Positive-disordered: experimentally-validated disordered regions from DisProt

The `DisProt` database [23] collects experimental annotations of intrinsically disordered regions. We worked with the June 2025 version of the database, and we selected the annotations for the 984 proteins having experimentally determined structures deposited in the Protein Data Bank. We then took the corresponding `AlphaFold2` predictions in order to have exact residue-wise mappings between the `DisProt` annotations and the structures, even for intrinsically disordered regions. We selected disordered regions of at least 10 consecutive amino acids according to the DSSP classification [25] (residues annotated with S, T, B, or -).

##### Negative-ordered: implausible motifs from our geometrical algorithm

We ran the algorithm detailed in Section 2.2 on the whole `AlphaFold-DB` human proteome and found a total of 20066 residues involved in implausible backbone conformations. Implausible regions (especially extremely twisted beta strands) are also occasionally found in PDB structures. This is to be expected, and only nuances the kNN classifier score.

##### Negative-disordered: disagreement between AlphaFold2 and AIUPred

The `AIUPred` algorithm [24] is a state-of-the-art disorder predictor that, although based on the transformer architecture, does not integrate PLM embeddings or `AlphaFold2`-derived information. `AIUPred` returns a score between 0 and 1 for each amino acid. A residue is likely part of an intrinsically disordered region when its score is ≥0.5. As a set of unconfirmed disordered regions predicted by `AlphaFold2`, we consider unstructured segments (DSSP codes S, T, B, or -) of more than 10 amino acids having a median `AIUPred` score of ≤ 0.3.

#### 2.3.3 Validation

The dataset `abstraqt` is trained on cannot be directly used for validation. Indeed, the defined labels are neither guaranteed to be exclusive nor exact: indeed, CATH structures contain residues with unusual backbone bends and twists, `AIUPred` has high but not perfect accuracy, and our algorithm only detects two main classes of hallucinations, others remain uncaught.

Since the extent of incomplete or erroneous labeling is unknown, we cannot rely on standard approaches like ROC/AUC to quantify the algorithm’s accuracy. Instead, we propose to quantify the orthogonality of `abstraqt` qith respect to pLDDT. We thus run `abstraqt` on the whole human proteome predicted by `AlphaFold2` and report cases where at least 10% of the residues have a High or Very High confidence (pLDDT ≥ 70) and `abstraqt` ≤ 0.3.

### 2.4 Quality assessment measures

We use the arity encoding to perform various analysis based on the following classical tools.

#### DisProt on proteins without experimentally determined structure

We worked with the June 2025 version of the `DisProt` database [23] which includes 413 disorder annotations for proteins that do not correspond to any PDB entry. The proteins come from 135 different species, spanning eukaryotes, prokaryotes and viruses. For each one of these sequences we compared the AFDB prediction and the `DisProt` annotation, where both were available, complete and unambiguous. For the remaining 273 proteins, we counted the percentage of residues that are described as holding secondary structure according to DSSP (H, I, G, and E labels) but are not annotated as disordered in `DisProt`.

#### AIUPred disorder prediction of whole proteomes

With the aim of analyzing the same problematics seen on `DisProt` but at the scale of a whole proteome, we computed `AIUPred` [24] predictions with standard parameters starting from the sequences of the `AlphaFold-DB` predictions, and counted the residues having `AIUPred` disorder score is ≥ 0.5 and DSSP label H, I, G, or E.

#### PROCHECK

We ran the `PROCHECK` v.3.5.4 [26] with resolution 1.00 A and collected the total G-factor of each prediction. The G-factor is a log-odds measure: a structure is considered and outlier when its G-factor is ≤ −1.

#### PhiSiCal-Checkup

We ran `PhiSiCal-Checkup` v.1.0 [27] which provides a per-residue score for a wide range of dihedral angle distributions. By default, outlier values are defined having a score of ≤ −4. For each protein, we counted the number of residues having at least one outlier score.

#### VoroMQA

`VoroMQA` [28] returns a per-structure normalized packing score. A score of *<* 0.5 determines a poor-quality structure in terms of packing. We ran `VoroMQA` on the whole human proteome and counted (per bin) the percentage of structures receiving a poor quality score.

### 2.5 Code availability

The code to compute arity maps is available within the the Structural Bioinformatics Library [29]. See: [AlphaFold analysis](https://sbl.inria.fr/doc/Alphafold_analysis-user-manual.html), as well as [SBL portal](https://sbl.inria.fr/), [Documentation](https://sbl.inria.fr/doc/), [Applications](https://sbl.inria.fr/applications/), [Installation guide](https://sbl.inria.fr/doc/sbl-installation-guide.html).

The data for rerunning analyses are made available from the SBL data website at [Data](https://sbl.inria.fr/data/AlphaFold-assessment).

## 3 Results

### 3.1 Arity is a faithful proxy of the 𝒲_2_ distance

The arity distribution quantifies the global packing properties, as seen from a comparison involving α and β folds, as well as proteins presenting unstructured regions or unconventional structures (Fig. 1). It is thus natural to use the Wasserstein 𝒲_2_ distance matrix between arity histograms of a set of proteins to organize the corresponding proteins into groups sharing the same local packing properties, effectively clustering well-folded, partially unfolded, and misfolded structures.

**Figure 1:**
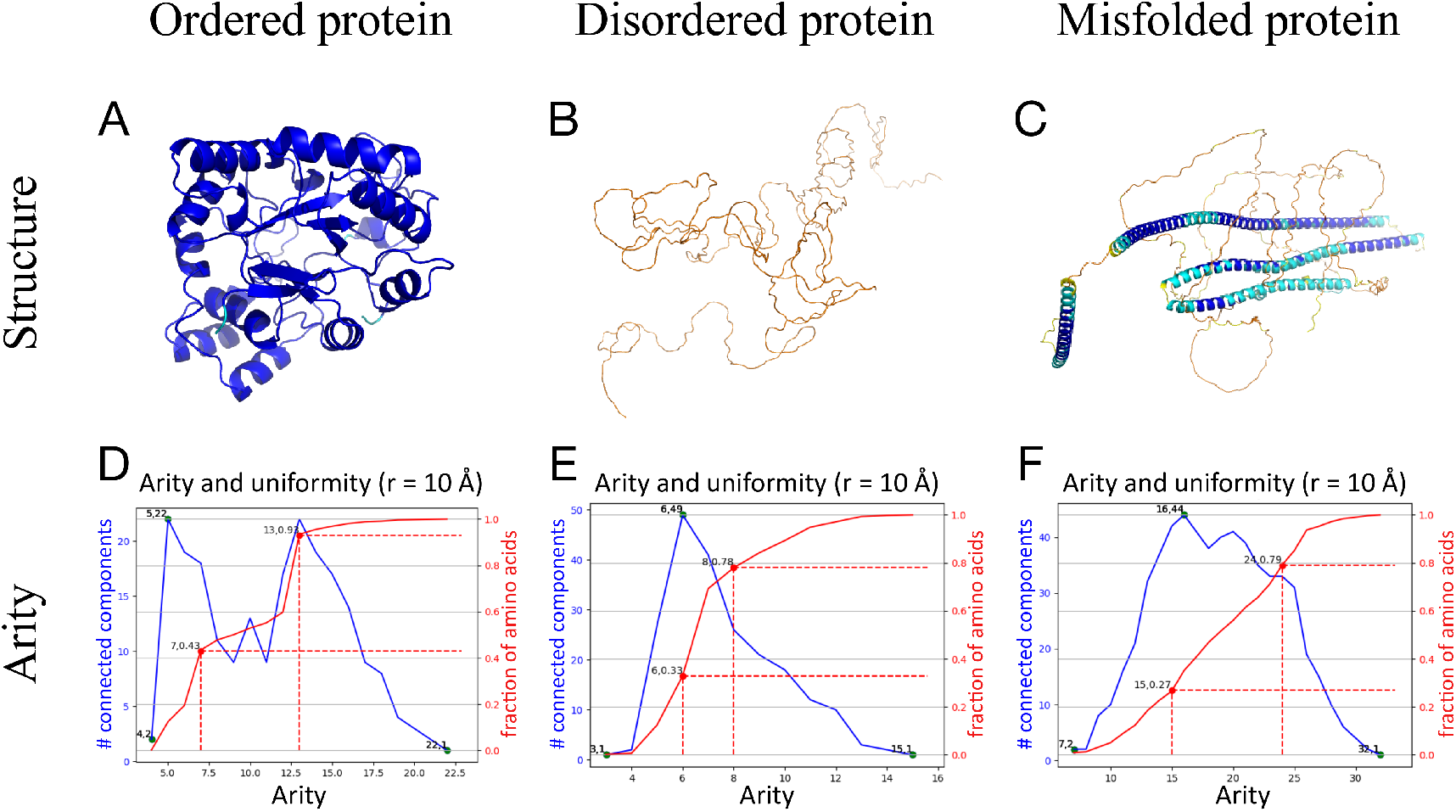
Arity of three prototypical examples corresponding to ordered, disordered and mixed proteins. **(A)** Protein1 (AF-P15121-F1-model_v4, *H. sapiens*), **(B)** Protein2 (AF-A0A0G2L439-F1-model_v4, *D. rerio*), **(C)** Protein3 (AF-Q9VQS4-F1-model_v4, *D. melanogaster*). **(D, E, F)** Red curve: arity CDF with the corresponding *q*_25_ and *q*_75_ values highlighted. Depending on the quantity and arrangement of structured regions, the arity shows one or more steep increases. Blue curve: number of connected components formed by the Union-Find algorithm upon inserting (into the sequence) a.a. by increasing arity.

We perform the Wasserstein analysis (Sec. 2.1.2). The first two principal components of such distance matrix explain respectively 45.2% and 4.2% of the variance of the dataset, with the third one accounting for only 0.8%. A two-dimensional PCA thus conveys most of the relevant information about global similarity.

We next investigate the relationship between this embedding and that associated with the arity map (Def. 2), conditioned to the two quantiles/percentiles *p*_1_ and *p*_2_ used to define it. For a given embedding (PCA or arity map), consider the *O*(*N* ^2^) distances in the embedding space. We compute the Spearman correlation between these two sets while varying the percentiles *p*_*i*_ ∈ {10, 20, 30, 40} and *p*_*j*_ ∈ {60, 70, 80, 90} . All Spearman values lie between 0.91 and 0.98, with the maximum at (30, 80) (Table 1). For simplicity, we select the percentiles *p*_*i*_ = 25 and *p*_*j*_ = 75. These are denoted *p*_25_ and *p*_75_ in the sequel, and *arity map* refers to Def. 2 with these values.

### 3.2 The arity map is consistent throughout species

We use the arity map to visualize structural characteristics of any large protein corpus at a glance. The x-axis 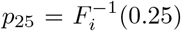 is geared towards loosely packed regions, while the y-axis 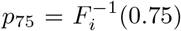 reflects tightly packed ones. For a given cell of the map, the distance to the diagonal *y* = *x* encodes the uniformity of the overall packing of the protein. For bins on the diagonal, a unique arity value suffices to move from a fraction of amino acids below 25% to a fraction above 75%, reflecting a highly uniform three-dimensional structure. This is never the case for native, non-trivial protein structures, which always present different quantiles *p*_25_ and *p*_75_ (Fig. 1(D-F)) due to their alternation between rigid and flexible regions.

For all proteomes, the densest region of the arity map locates in a short diagonal strip centered in [15, 21] referred by *native sector*, or NS, in the following, which corresponds to well-folded proteins (Fig. 2(A)). Other densely populated regions are typically found in the wider [5, 5] × [15, 15] sector (*heterogeneous sector*, or HS), but their occurrence and exact location largely depends on the species and the quality of predictions. For example, *P. falciparum* exhibits a continuum between the two sectors, whereas *E. coli* only present high density in the native sector. The case of *H. sapiens* is also remarkable as it presents a particularly populated bin at 7 × 13 in addition to the more typical denser region at very low arity values.

**Figure 2:**
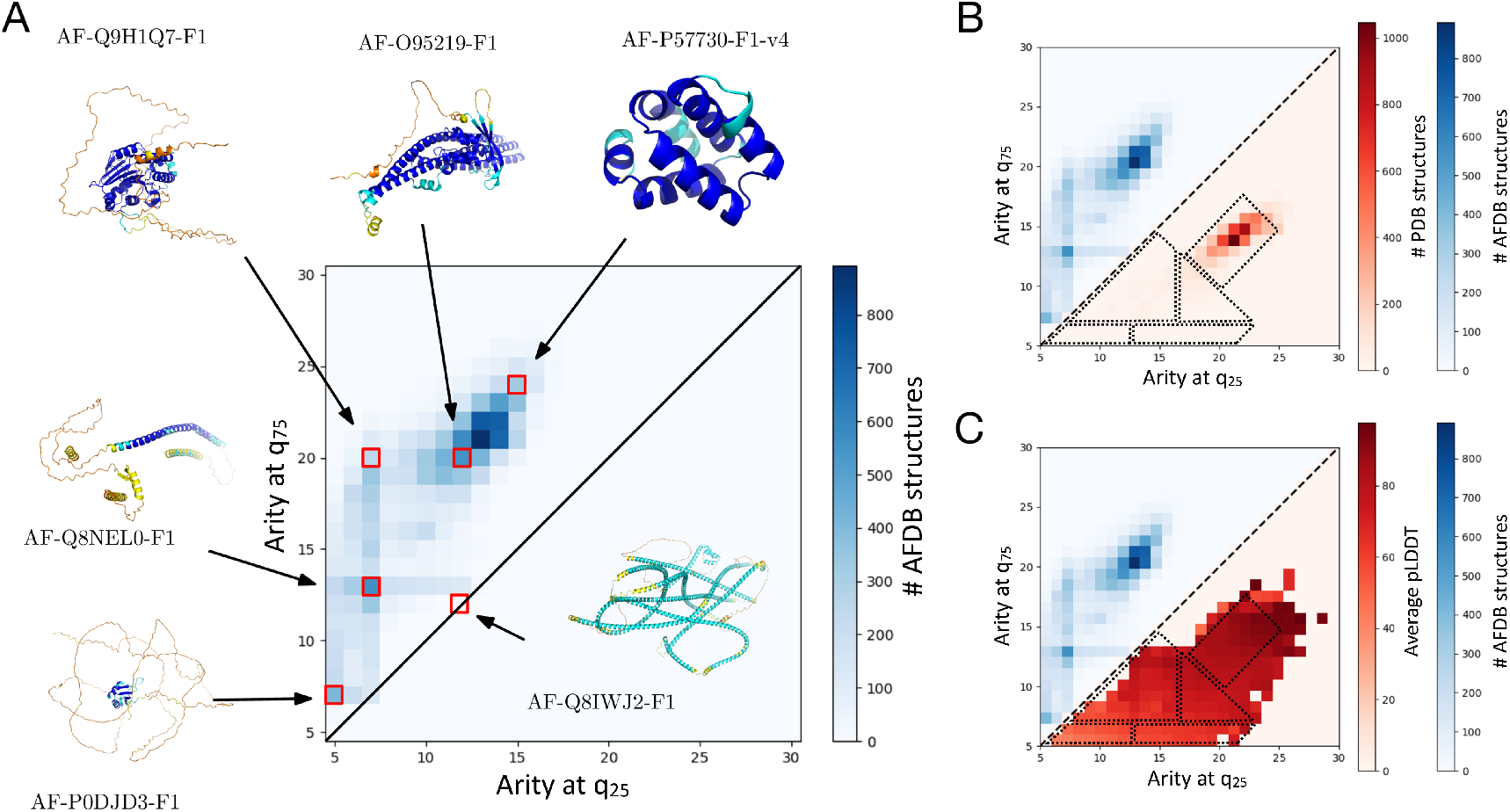
Arity map of the whole *H. sapiens* genome. The upper triangle of the arity map always shows the protein structure count for each (*q*_25_, *q*_75_) arity signature. **(A)** Each bin corresponds to all `AlphaFold2` predictions harboring a given arity signature. For some key signature, a representative protein structure is shown. The largest and highest density corresponds to well-folded structures (that can also include flexible or intrinsically disordered regions), the bottom-most region of the arity map corresponds to intrinsically disordered proteins with little or no structure, and the region around the density peak in [7 × 13] is enriched in partially folded and misfolded structures. **(B)** Lower triangle – mirrored with respect to the upper triangle: protein count from the PDB. Well-formed, ordered proteins only populate the (15,20) peak of the arity plot, confirming that the other two peaks visible in the arity plot of the `AlphaFold-DB` predictions must represent disordered structures or inaccurate predictions. Dashed lines are consensus boundaries (see Discussion). **(C)** Lower triangle: average pLDDT per bin. Arity and pLDDT are strongly correlated: the higher the first and last quartile, the higher the average pLDDT. Arity is thus a good estimator of structural complexity. Dashed lines are consensus boundaries (see Discussion).

In the next three sections, we present three example analyses on the ∼ 23, 000 protein structures predicted by `AlphaFold2` and deposited in·`AlphaFold-DB`. The results we reach can easily be validated by conducting the same calculations on other genomes from `AlphaFold-DB`, using the software accompanying this work, available within the Structural Bioinformatics Library [29], see [AlphaFold analysis](https://sbl.inria.fr/doc/Alphafold_analysis-user-manual.html).

### 3.3 Folded, unfolded, and misfolded structures group in different regions of the arity map

In order to confirm that well-folded proteins are mapped in the native sector, we compared the arity maps of the AF2-predicted human proteome and the set of all human protein structures deposited on the PDB. As expected, the PDB structures only cluster in the native sector (Fig. 2(B)). Only 32 structures out of the 73169 deposited on the PDB populate the 7 × 13 bin commented in the previous section. All of them are NMR structures of microproteins counting ≤73 a.a., with partially unfolded structure. The arity map can thus cluster well-folded protein structures. In passing, we note that pLDDT is correlated with arity: structures with a high (low) arity signature will also have a high (low) average pLDDT value (Fig. 2(C)), confirming that the arity is also a good proxy for prediction robustness. Yet, pLDDT is not able to show the finer details of the structural repartition that emerges from the arity map.

Proteomes of all species also present considerable percentages of proteins with intrinsically disordered regions IDR, which in eukaryotes can often sum up to 20-30% [30]. `AlphaFold2` has been found to reliably predict intrinsic disorder, describing IDRs as unwound, maximally extended regions. Nonetheless, artifacts like the prediction for AF-Q8IWJ2-F1 (Fig. 2(A)) also present looser packing. Is the arity map able to separate correctly-predicted intrinsically disordered regions and proteins from artifacts?

In order to quantitatively characterize the arity map density population, we mapped the domains described in the recent TED database [3] on the human proteome (Fig. 3). TED domains are grouped in three quality classes, depending on whether consensus has been found among three different approaches for recognizing structural domains.

**Figure 3:**
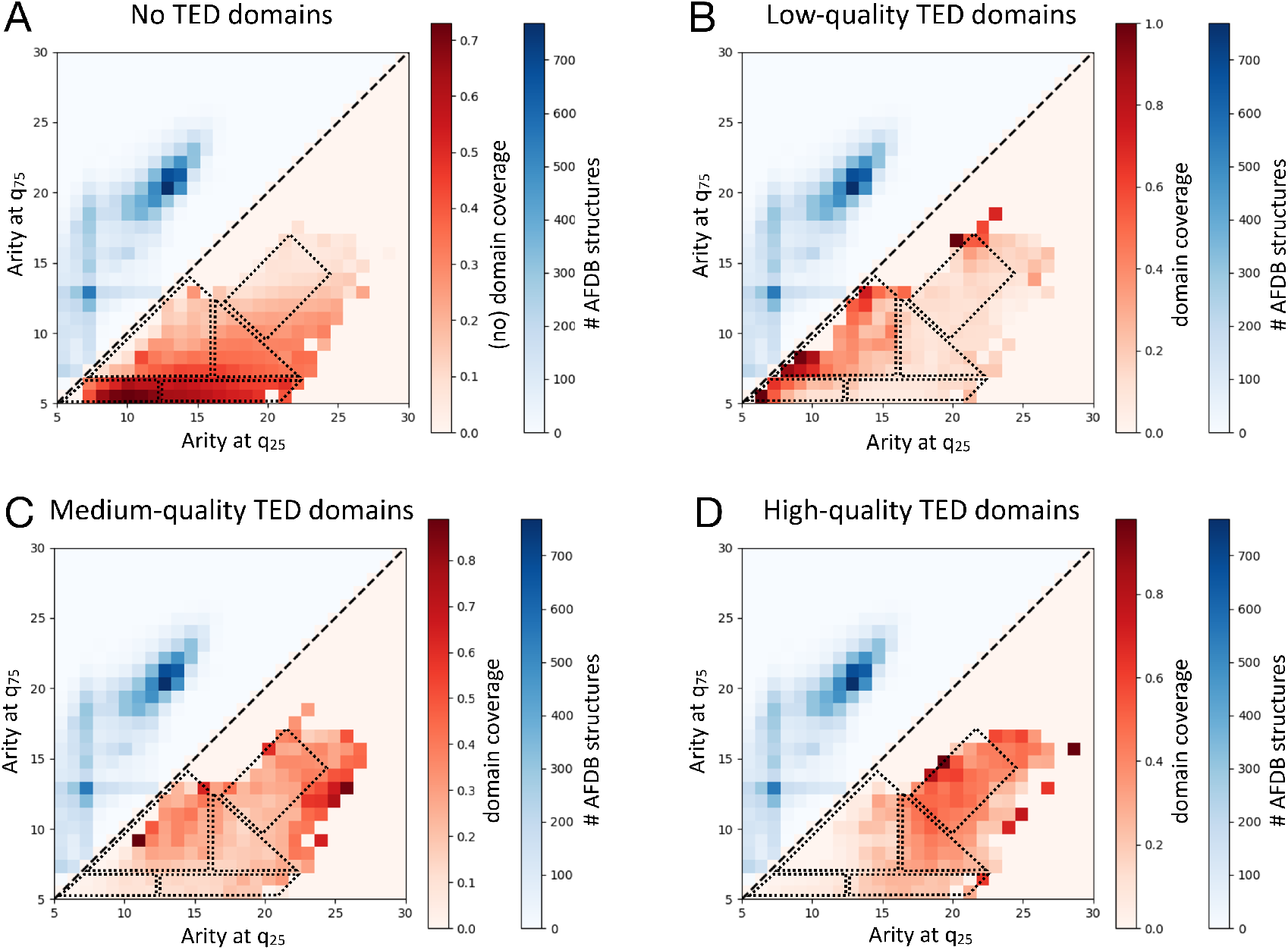
Arity signatures of TED domains divided by quality. For each protein, the fractions of its a.a. with a TED label are computed – these numbers add up to one. A cell of the arity map displays the average value of the fraction corresponding to a given label. The arity map distinguishes between structures having **A)** no, **B)** low-quality, **C)** medium-quality, and **D)** high-quality TED domains.

For each protein, the fraction of amino acids assigned to each TED class is computed, with the fractions summing to one. Each cell of the arity map represents the average fraction associated with a given class.

As expected, high-quality (consensual) domains are enriched in the native sector, whereas medium- and low-quality domains are predominantly present in the heterogeneous sector, albeit in two different locations. The heterogeneous sector thus splits in three different sub-locations, one enriched in low-quality domains, one in medium-quality domains, and another where domain identification is scarce.

### 3.4 Unrecognized intrinsically disordered regions tend to be predicted as long alpha helices

Among the most recurrent `AlphaFold2` hallucinations there is the tendency to predict secondary structure in intrinsically disordered regions (IDRs). We present two analyses, one based on experimental data and one using a state-of-the-art disorder predictor.

#### DisProt experimentally validated IDRs

We first consider the `DisProt` database [23], which collects disordered regions from more than 1300 target protein sequences. Many of the benchmarked sequences correspond to proteins of known structure, present in the PDB. For this test, we focus on the 273 targets lacking structural data and having a predicted structure on `AlphaFold-DB`. The proteins are issued from 135 different species, and each residue is annotated as intrinsically disordered or not. We compute the fraction of experimentally confirmed ID residues that are predicted by `AlphaFold2` as having secondary structure– we term these *False Negative disordered residues*. For arity signatures with little difference between the two quantiles (e.g. (10, 12), (14, 15)), there can be as much as 80% of hallucinated secondary structures, mostly consisting in long alpha helices (Fig. 4(A)).

**Figure 4:**
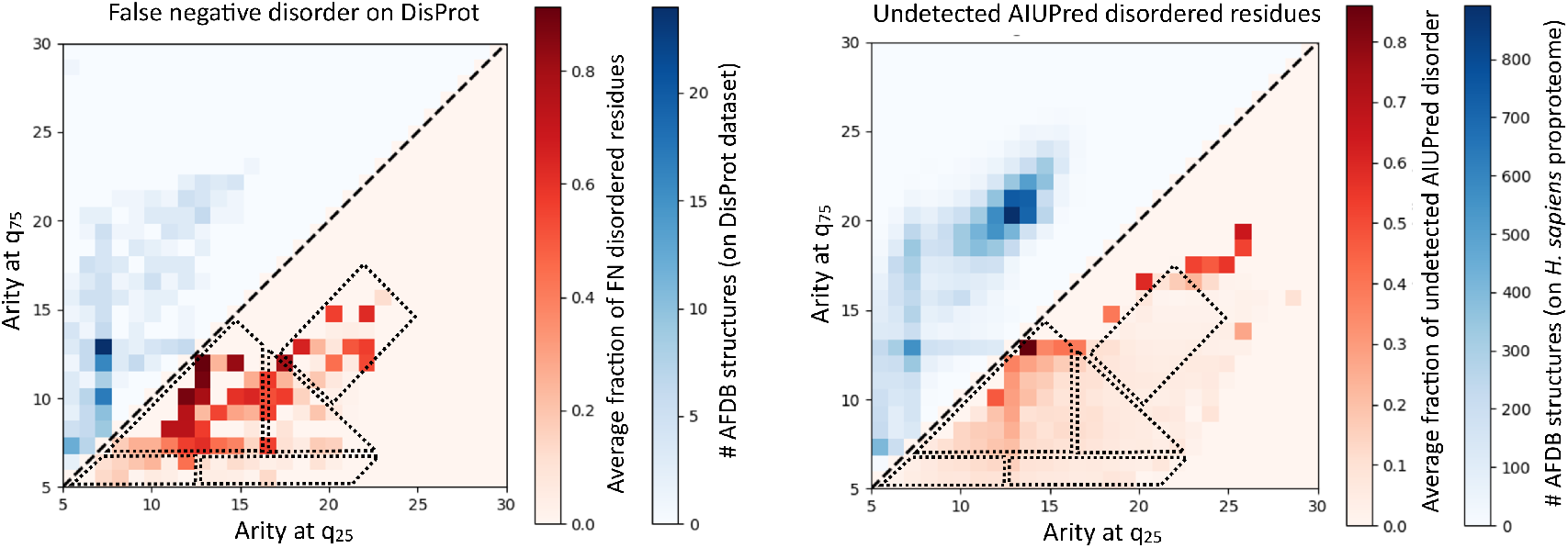
H. Sapiens: arity map groups structures with unrecognized intrinsically disordered regions. **(A)** `DisProt` analysis on 273 target proteins from 135 different species lacking structural data. The lower triangle of the arity map shows the fraction of amino acids with secondary structure but annotated as disordered in `DisProt`: these are False Negative disordered residues in `AlphaFold2`’s predictions. Dashed lines are consensus boundaries (see Discussion). **(B)** `AIUPred` analysis on the human proteome. The lower triangle reports the average number of residues with secondary structure but also `AIUPred` *>* 0.5. The enriched area is a subset of the one highlighted in `DisProt` corresponding to its most populated bins, confirming that the arity map groups proteins where `AlphaFold2` has hallucinated ordered conformations out of intrinsically disordered regions. Dashed lines are consensus boundaries (see Discussion).

#### Analysis using AIUPred

`AlphaFold2` predictions have been found to generally agree with state-of-the-art disorder predictors, among which `AIUPred` [24], a recent neural-network-based implementation of IUPred trained on IUPred predictions. `AIUPred` predicts a residue as disordered when the `AIUPred` score is *>* 0.50. We therefore use the arity map to study the (lack of) homogeneity between the `AIUPred` annotation and the `AlphaFold2` prediction. The same region that is enriched in mispredicted `DisProt` IDRs also contains the majority of incoherent annotations between `AlphaFold2` and `AIUPred` (Fig. 4(B). The enrichment signal in the region around 17 × 25 is instead an artifact given by the extremely low number of structures per bin – usually 1-2 against the 30-200 of the main region.

### 3.5 AF2 misfolds are not caught by standard quality checks

To challenge the discerning power of the arity map, we also use standard stereochemical quality checks such as `PROCHECK` [26] and `PhiSiCal-Checkup` [27] to spot AF2 misfold predictions. Both algorithms flag a very small number of predictions as problematic, including physically implausible structures.

`PhiSiCal-Checkup` evaluates the stereochemistry of each residue by calculating a log-odd score for several joint dihedral distributions. Following the default definition, we classified as outliers residues having at least one log-odd score of -4 or less. We then calculate the average fraction of outlier residues per prediction in each bin of the arity map, and we see that it correlates with predicted disorder (Fig. 5(A)).

**Figure 5:**
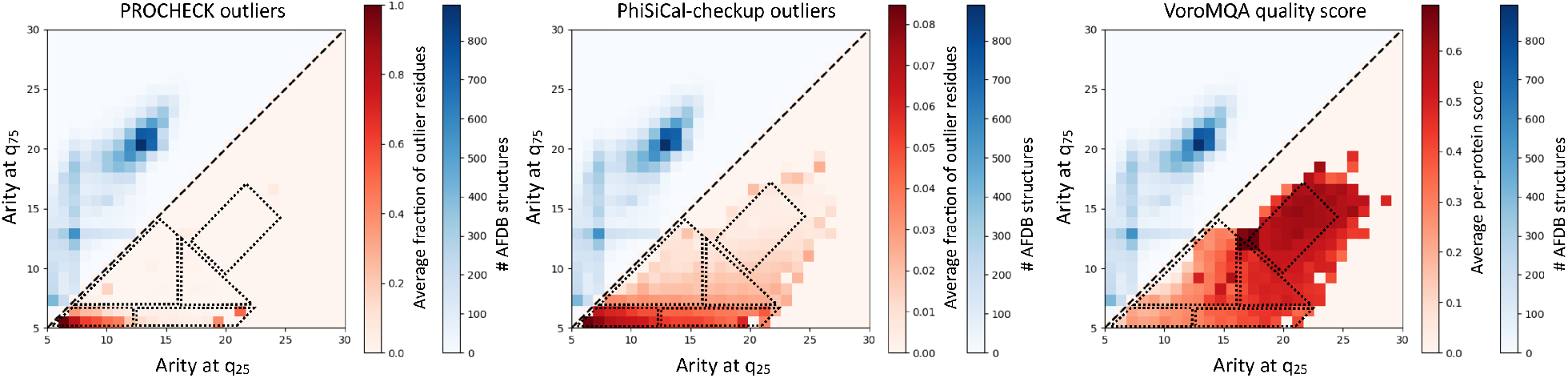
Comparison with structure quality assessment measures. We plot on the arity map the results from two structure quality assessment algorithms. Dashed lines are consensus boundaries (see Discussion).**(A)** For `PhiSiCal-Checkup`, we plot the average percentage of residues per predicted structure having at least one rotamer score of -4 or less. **(B)** For `PROCHECK`, we plot the percentage of structures having a global G-factor of -1 or less. **(C)** For `VoroMQA`, we plot the average `VoroMQA` score of each structure.

`PROCHECK` evaluates the stereochemical quality of protein structures by means of G-factors, log-odds scoring functions that quantify how unusual certain stereochemical parameters are. A G-factor of -1 or less corresponds to a problematic structure. Like in the case of `PhiSiCal-Checkup`, highlighted structures concentrate in the unfolded part of the arity map, disregarding misfolds (Fig. 5(B)).

If stereochemistry-based scoring function cannot grasp `AlphaFold2` misfolds, packing-based methods could. A widely used technique in this respect is Voronoi tessellation. `Voronoia` is a widely used Voronoi tessellation package that also includes a quality assessment method called `VoroMQA`. The method returns a normalized packing score based on the deviation of the volume of the Voronoi cells centered on each residue from the distribution observed for experimentally determined, well-folded structures. A score of *<* 0.5 determines a poor-quality structure in terms of packing. We ran `VoroMQA` on the whole human proteome and counted the percentage of poor-quality packing score per arity map bin: the only region containing good-quality structures coincides with the Native Sector (Fig. 5(C)). `VoroMQA` is thus unable to make the difference between AlphaFold’s representation of truly disordered regions and loosely-packed misfolds.

The arity map has thus more discerning power than standard structure quality assessment measures, whether they are based on dihedral angle or packing distributions.

### 3.6 abstraqt: a residue-wise structure quality assessment measure

We have so far considered local packing analysis for simple and efficient qualitative visualization of large sets of predicted protein structures. The same instrument can be used to compute a per-residue local structural quality measure analogous to pLDDT – Sec. 2.3.

The kNN based regressor `abstraqt` (Arity-Based STRuctural Arrangement Quality assessmenT) delivers a per-residue score on a scale from 0 (worst) to 100 (best). While agreeing with pLDDT on well-packed regions, `abstraqt` is able to recognize unphysical secondary structures such as long curved alpha helices or unstable beta strand conformations (Fig. 6(B-C) and (D-E) respectively). Disordered regions are scored depending on their structural context, and both globular and non-globular crystallographic structures receive uniform, high scores (Fig. 6(F-G)). When evaluating the whole *H. sapiens* `AlphaFold-DB` proteome, we find that 3711 / 1120 / 480 / 106 / 4 out of 23393 (15.8% / 4.8% / 2.1% / 0.5% / 0.02%) structure predictions contain at least 10% / 30% / 50% / 70% / 90% residues with pLDDT ≥ 70 and `abstraqt` ≤0.3. The residues showing score disagreement majoritarily compose long, bent helices (3656 records), followed by unstructured regions (1534 records) and beta strands (361 records). The enrichment in low-score residues (`abstraqt` ≤0.3) over the arity map (Fig. 6(A)) highlights the same areas found in previous sections, notably the disagreement between `AlphaFold2` and `DisProt`/`AIUPred` and the Low-quality TED domains.

**Figure 6:**
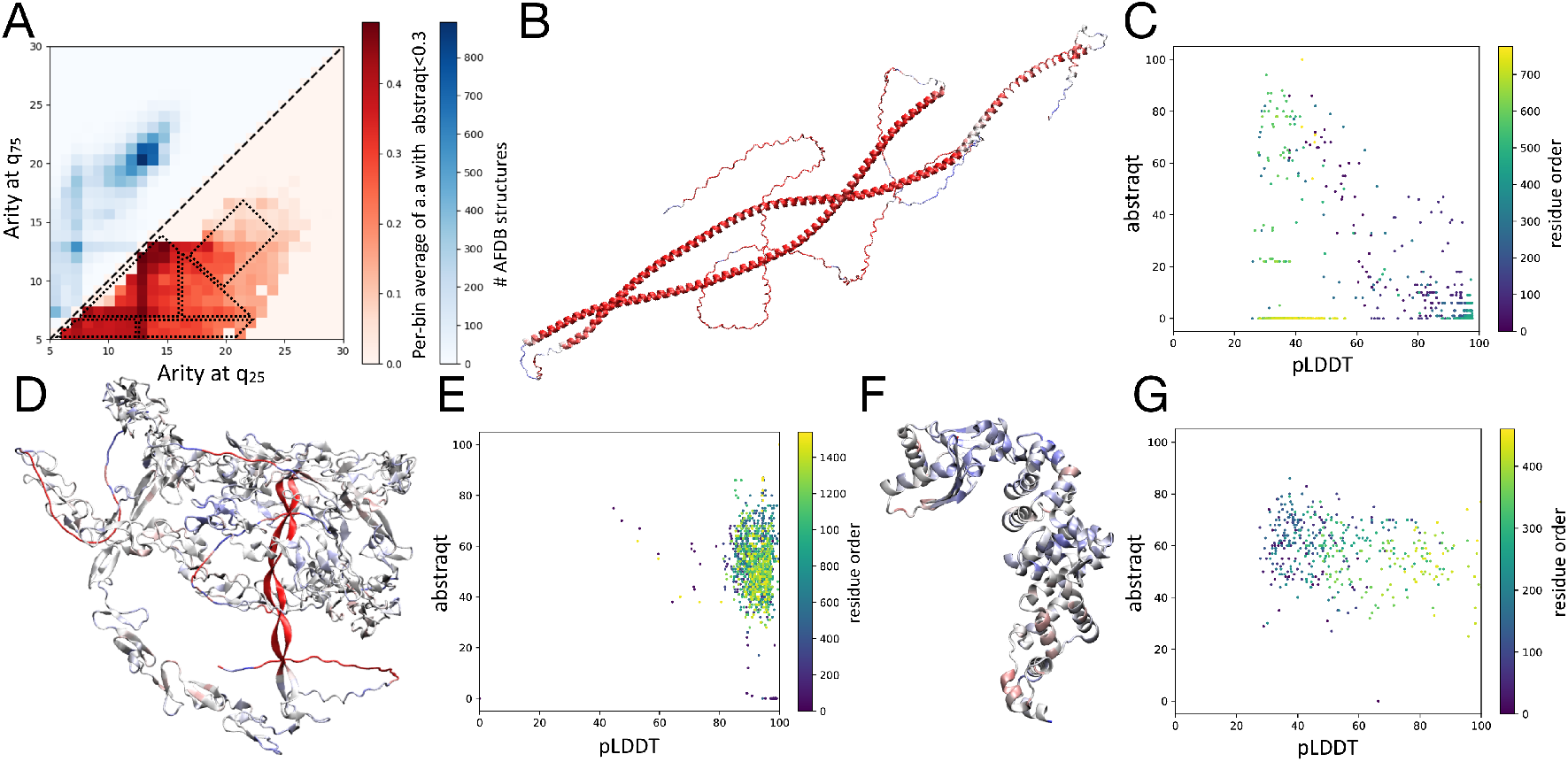
The abstraqt per-residue score on pathological and control structures. **(A)** Per-bin average of the fraction of residues having `abstraqt` score ≤0.3. Dashed lines are consensus boundaries (see Discussion).**(B, D)** Prediction of two proteins (`AF-A0A2R8YE69-F1-model_v4` and `AF-O75095-F1-model_v4` respectively) displaying large implausible motifs, highlighted by a very low `abstraqt` per-residue score (color code: red - 0, grey - 0.5, blue - 1). **(C, E)** For the two examples, the correlation between `abstraqt` and pLDDT is shown. Each point represents a residue in the prediction, and is colored according to its position in the polypeptide chain. The first example shows a marked anticorrelation between the two scores (the long helices have a high pLDDT score), the second shows how `abstraqt` distinguishes between plausible and implausible regions whereas pLDDT is very high everywhere. **(F, G)** Control case (PDB code 9NS2) showing that crystallographic non-globular proteins are associated with overall high and uniform `abstraqt` values.

## 4 Discussion

We introduce a packing-based framework to assess `AlphaFold2` predictions at proteome and residue scales. Each protein is represented by the histogram of residue neighbor counts– at inclusion radius 10 Å, and packing similarity between proteins is measured using the Wasserstein 𝒲_2_ distance between normalized histograms. Neighbor count distributions are sensitive to structural compactness, while the Wasserstein distance provides a statistically meaningful comparison of distributions. A statistical analysis based on the Wasserstein PCA shows that the space of arity distribution is well captured by two principal vectors–explaining ∼ 50% of the variance. This observation is leveraged by defining the arity map, a 2D embedding of proteins based on the quantiles associated to the 25th and 75th percentiles of its cumulative distribution. The resulting two-dimensional histogram therefore offers a lightweight and interpretable approximation of proteome-wide packing structure. Because it relies only on neighbor count distributions, the method generalizes across species and prediction systems without recalibration.

The only part of the arity map that is populated by structures taken from the PDB is a diagonal rectangular area at high (*q*_25_, *q*_75_) he call the *native sector* (cfr. Figure 2B). The remaining area showing non-negligible density is called *heterogeneous sector*.

The inability of pLDDT to detect implausible predictions is especially important. Although higher pLDDT values generally coincide with the native sector, pLDDT often assigns high confidence to physically unrealistic structures if their local geometry is internally consistent. This mirrors the behavior observed for adversarial sequences, where a few substitutions can drastically alter the predicted fold without reducing confidence. The arity map captures these cases because it reflects anomalous global packing rather than local geometry alone. For example, a long curved helix creates globally atypical packing environments despite having ordinary backbone dihedrals.

The relationship between arity and intrinsic disorder is also informative. `AlphaFold2` typically represents intrinsically disordered regions as extended and loosely packed chains, which occupy the low-arity region of the map. Nonetheless, there exists a sub-region of the arity map containing proteins where `AlphaFold2` predicts long alpha helices in regions classified as disordered by `DisProt` and `AIUPred`. Importantly, this sub-region is consistent between `DisProt` and `AIUPred` and spatially separated in the arity map, indicating that arity alone can distinguish plausible disorder from hallucinated structure. This distinction is practically relevant because filtering solely by pLDDT may retain structured artifacts in disorder analyses.

Standard quality metrics do not discriminate hallucinations. `PROCHECK` and `PhiSiCal-Checkup`, both based on dihedral angle distributions, mainly identify outliers in the disordered regions of the arity map and fail to detect implausible structures. Hallucinated helices and beta solenoids may retain acceptable local geometry while remaining globally unrealistic. `VoroMQA`, despite evaluating packing through Voronoi tessellation, also performs poorly by assigning low scores broadly across the Heterogeneous Sector without separating disorder from misfolding. The arity map therefore provides higher discriminatory power at lower computational cost.

Combining these evidences, we propose a qualitative yet meaningful and informative partition of heterogeneous sector into regions corresponding to different enrichments (Fig. 7):

**Figure 7:**
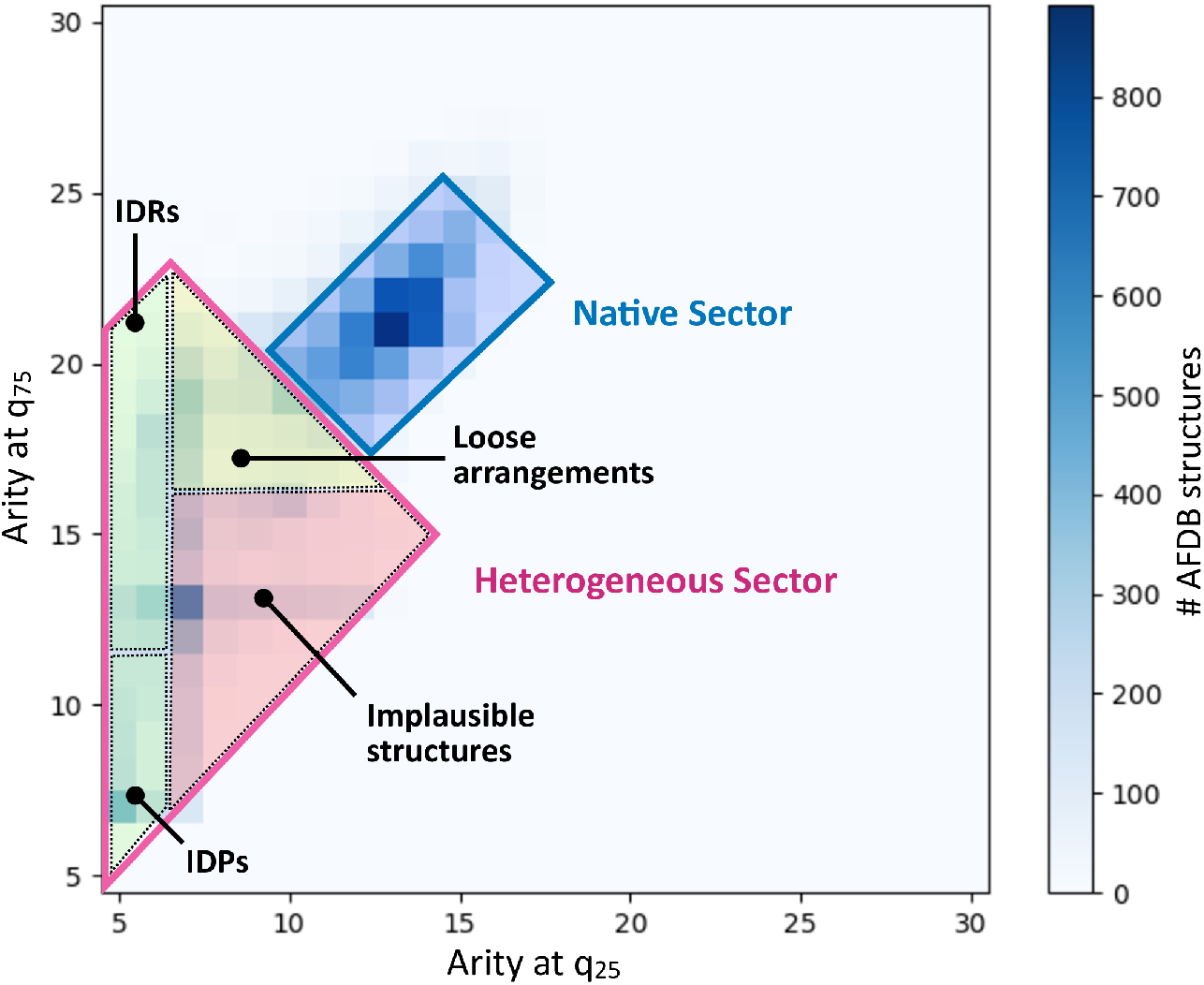
Arity map sectors. We propose a partition of the arity map into meaningful regions corresponding to different enrichments. The native sector contains most well-folded, compact structures, and is the only one populated by PDB structures. The heterogeneous sector contains structures that do not resemble crystallographic structures, and can be divided into four regions: the IDP region comprises completely unfolded structures with no TED annotations. The IDR region contains structures with large intrinsically disordered regions, validated but disorder predictors. The Implausible structures region is enriched in Low quality TED domains and cases where `AlphaFold2` disagrees with `DisProt` and `AIUPred` over disorder. Finally, the Loose arrangements region contains proteins enriched in High-quality TED domains, but whose arrangement is not as compact as an experimentally validated structure.

- The *IDP region* (low *q*_25_ and *q*_75_) comprises completely unfolded structures with no TED annotations (Fig. 3(A)). Disordered structures are validated by `DisProt` experimental evidence or `AIUPred` independent predictions (cfr. Fig. 4). Quality assessment scores are generally low because of the random orientation of backbone angles and the low degree of compactness (Fig. 5)
- The *IDR region* (low *q*_25_, high *q*_75_) contains intrinsically disordered regions. An enrichment in structures with no TED domains is still present, but whereas there are annotated TED domains, they are of High quality (Fig. 3(A) and (D)). Quality assessment scores are still low due to the presence of disordered regions
- The *implausible structures region*·is characterized by a marked enrichment in Low quality TED domains (Fig. 3(B)), and in regions that appear structured in `AlphaFold2` but that correspond to `DisProt` disorder annotations or `AIUPred` disorder predictions (Fig. 4).
- Finally, the *loose arrangements region* is enriched in High-quality TED domains and does not present a high percentage of disorder. `AIUPred` does not predict a high content of disordered regions and quality assessment functions flag a negligible number of amino acids. Yet, only about 200 PDB structures out of 73169 have the corresponding arity signatures. Upon visual inspection, it seems the majority of these structures are composed of or contain meaningful domains, but their arrangement is not compact.

At the local *i*.*e*. amino acid level, the ability of our scoring function `abstraqt` to recognize unphysical secondary structures (long curved alpha helices, unstable beta strand conformations) and unlikely structured/disordered regions is especially interesting to separate the wheat from the chaff and still enjoy reliable regions in a model. When queried on the whole human proteome, `abstraqt` retrieves the same regions described for the arity map (Fig. 6(A)).

We note in passing that although the `abstraqt` training set is diverse, it cannot yet cover all structural contexts present in the expanding `AlphaFold-DB`. While the non-parametric Ball tree approach should improve as more data become available, systematic validation on newly solved structures is still required.

Overall, the arity map provides a lightweight and interpretable complement to existing AlphaFold2 quality metrics for large-scale proteome studies. Its computational cost is minimal and requires only predicted PDB structures as input. The `abstraqt` score extends this analysis to residue resolution without requiring additional sequence alignments, evolutionary data, or neural network inference. Together, these tools offer a practical framework for improving automated curation of `AlphaFold-DB` predictions, with applications in structural genomics, drug discovery, and the analysis of intrinsically disordered proteomes.

## Acknowledgments

This work has been supported by the French government, through the 3IA Côte d’Azur Investments (ANR-19-P3IA-0002), and the ANR project Innuendo (ANR-23-CE45-0019).

The authors wish to thank the reviewers for their accurate evaluation and constructive recommendations.

