## Supporting Information for "Fold or flop: quality assessment of AlphaFold2 predictions on whole proteomes"

### S1 Supporting information

###### S1.0.1 Inclusion radius

There is no consensus in the literature on a $C_\alpha - -C_\alpha$ distance threshold for defining neighboring residues, which is typically chosen in the range 8–14 Å depending on the application. We calculate the two-way trustworthiness between k-neighborhoods ($k = \{10, 20, 50, 100\}$) in the PCA and arity map spaces and find values in the range $[0.95, 0.97]$ for any radius between 6 Å (capturing only secondary structure contacts) and 14 Å (capturing the second residue shell). As a compromise between informativeness and computational cost, we fix the neighborhood distance threshold at 10 Å; all quartile values and references to the arity map hereafter use this threshold.

###### S1.0.2 Wasserstein PCA and arity map embedding

| $q_1$ | $q_2$ | Spearman coeff. |
|---|---|---|
| 10 | 60 | 0.919 |
| 10 | 70 | 0.944 |
| 10 | 80 | 0.943 |
| 10 | 90 | 0.905 |
| 20 | 60 | 0.930 |
| 20 | 70 | 0.960 |
| 20 | 80 | 0.970 |
| 20 | 90 | 0.948 |
| 30 | 60 | 0.937 |
| 30 | 70 | 0.967 |
| 30 | 80 | **0.980** |
| 30 | 90 | 0.967 |
| 40 | 60 | 0.936 |
| 40 | 70 | 0.965 |
| 40 | 80 | 0.978 |
| 40 | 90 | 0.970 |

Table S1: **Grid search over the percent values of the arity map quantiles.** We calculate the Spearman correlation coefficient between the 2-dimensional PCA of $\mathcal{W}_2$ distances and the arity map, a 2D-histogram defined over two quantiles corresponding to the $i$ and $j$ percentages. The best correlation coefficient between the two 2D dimensionality reductions corresponds to the (30, 80) percent pair, thus selecting quantiles very close to the first and third quartile.

###### S1.0.3 Arity distribution for typical folds

#### S1.1 pLDDT statistics per organism

For the reference, this section provides simple analysis on pLDDT values.

Let us recall the following intervals for pLDDT values: very high: $0.9 \leq$ pLDDT; high: $0.7 \leq$ pLDDT $<$ 0.9; low: $0.5 \leq$ pLDDT $< 0.7$; very low pLDDT $< 0.5$.

While the distribution of pLDDT values has been studied in selected cases, in particular for H. Sapiens in the context of conditionally folding proteins, we perform instead a systematic study on all genomes. We process all predictions (PDB files from `AlphaFold-DB`) of a given organism, collecting, per prediction, all/the median/the mean pLDDT value(s). These values yield three distributions, namely that for (i) all pLDDT values of all a.a. of all proteins, (ii) all median pLDDT values, and (iii) and all mean pLDDT values (Table S2).

Per genome statistics show that the fraction of a.a. with pLDDT below 0.7 lies in the range 0.078-0.408 and is rather consistent for model organisms (Table S2). However, the inspection of other organisms shows major discrepancies, with a value as low as 0.086 for *P. aeruginosa* and as high as 0.584 for *P. falciparum* (Table S2, Table S4, Table S5)

Selected illustrations are provided below.

| genome / pLDDT | 50 | 70 | 90 |
|---|---|---|---|
| AThaliana/all | 0.204 | 0.315 | 0.550 |
| CAlbicans/all | 0.213 | 0.312 | 0.570 |
| CElegans/all | 0.202 | 0.323 | 0.597 |
| DDiscoideum/all | **0.288** | 0.423 | **0.681** |
| DMelanogaster/all | 0.281 | 0.387 | 0.623 |
| DRerio/all | 0.246 | 0.343 | 0.593 |
| EColi/all | **0.029** | **0.078** | 0.275 |
| GMax/all | 0.217 | 0.338 | 0.577 |
| HSapiens/all | 0.284 | 0.382 | 0.666 |
| MJannaschii/all | 0.035 | 0.084 | **0.264** |
| MMusculus/all | 0.255 | 0.352 | 0.597 |
| OryzaSativa/all | 0.257 | **0.408** | 0.623 |
| RattusNorvegicus/all | 0.253 | 0.351 | 0.596 |
| SCerevisiae/all | 0.213 | 0.314 | 0.582 |
| SPombe/all | 0.187 | 0.289 | 0.558 |
| ZeaMays/all | 0.260 | 0.394 | 0.635 |
| SAureus/all | 0.045 | 0.096 | 0.295 |
| HPylori/all | 0.053 | 0.123 | 0.347 |
| MTuberculosis/all | 0.069 | 0.133 | 0.322 |
| Aeruginosa/all | **0.036** | **0.086** | **0.278** |
| PFalciparum/all | **0.460** | **0.584** | **0.804** |

Table S2: **Cumulated distribution function for pLDDT values of all a.a. of all proteins in a genome.** Top: model genomes. Bottom: five *global health* proteomes. Min and max values per column stressed in bold.

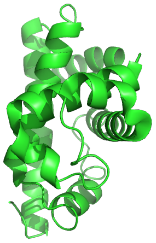

Fold $\alpha$

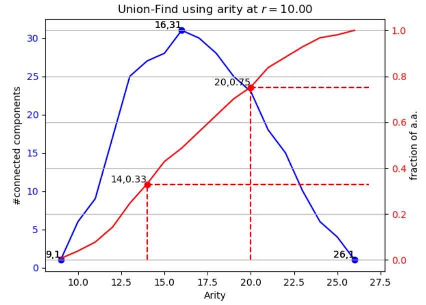

Globin, 154 a.a.

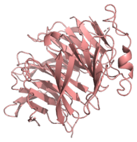

Fold $\beta$

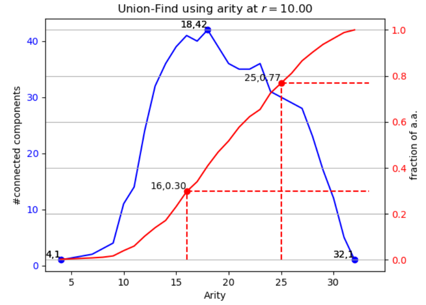

Propeller, 350 a.a.

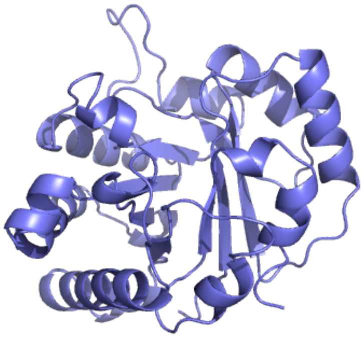

Fold $\alpha/\beta$

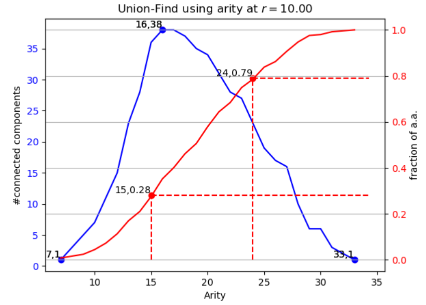

TIM barrel, 247 a.a.

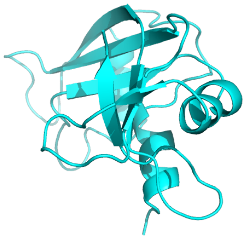

Fold $\alpha + \beta$

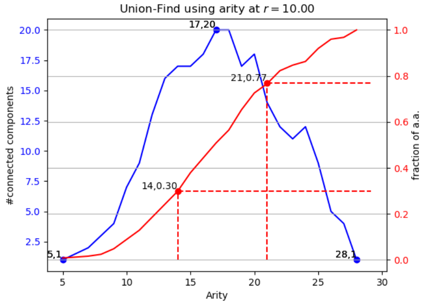

Ribonuclease, 124 a.a.

Figure S1: **Arity distribution and arity signature for prototypical folds.** Fold $\alpha$: 101m-globin-alpha; 154 amino acids. Quantile-arity signature [(0.25, 14), (0.5, 17), (0.75, 20)] Fold $\beta$: 1erj-propeller-beta; 350 amino acids. Quantile-arity signature [(0.25, 16), (0.5, 20), (0.75, 25)] Fold $\alpha/\beta$: 8tim-TIMbarrel-alphaSlashBeta; 247 amino acids. Quantile-arity signature [(0.25, 15), (0.5, 19), (0.75, 24)] Fold $\alpha + \beta$: 1a5p-ribonuclease-alphaPlusBeta; 124 amino acids. Quantile-arity signature [(0.25, 14), (0.5, 17), (0.75, 21)]

##### S1.1.1 HSapiens

As reported in Figure S2 and Table S3:

- Worst cases, top 3: ['File HSapiens/AF-Q96MU5-F1-model_v4.pdb has 243 a.a.; pLDDT min/med/max: 19.88 25.35 38.27', 'File HSapiens/AF-Q6ZR03-F1-model_v4.pdb has 302 a.a.; pLDDT min/med/max: 21.04 26.27 46.20', 'File HSapiens/AF-Q96M85-F1-model_v4.pdb has 177 a.a.; pLDDT min/med/max: 20.02 26.32 42.34']
- Median cases, 3 of them: ['File HSapiens/AF-P56159-F1-model_v4.pdb has 465 a.a.; pLDDT min/med/max: 25.40 86.54 98.58', 'File HSapiens/AF-Q96N96-F1-model_v4.pdb has 652 a.a.; pLDDT min/med/max: 24.37 86.54 98.41', 'File HSapiens/AF-Q9H009-F1-model_v4.pdb has 215 a.a.; pLDDT min/med/max: 36.28 86.54 95.84']
- Top cases, top 3: ['File HSapiens/AF-Q99497-F1-model_v4.pdb has 189 a.a.; pLDDT min/med/max: 67.04 98.81 98.97', 'File HSapiens/AF-P00352-F1-model_v4.pdb has 501 a.a.; pLDDT min/med/max: 28.66 98.82 98.98', 'File HSapiens/AF-P21549-F1-model_v4.pdb has 392 a.a.; pLDDT min/med/max: 58.71 98.83 98.98']

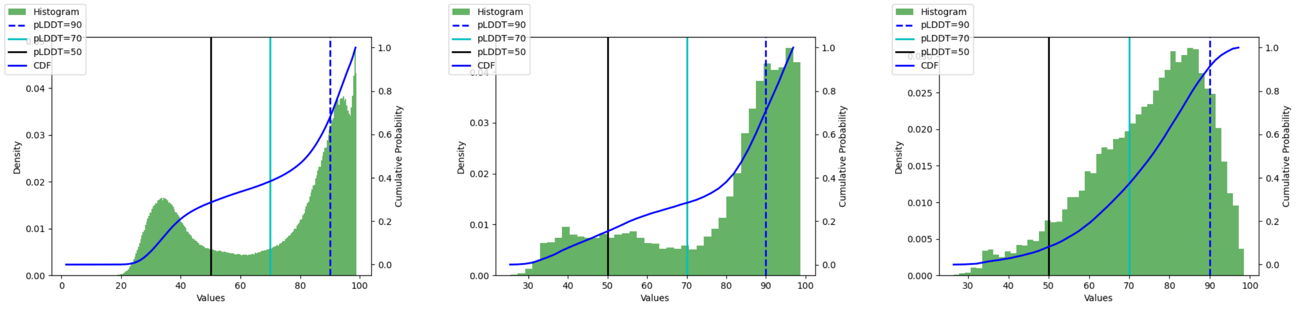

pLDDT all    pLDDT median    pLDDT mean

Figure S2: **HSapiens: pLDDT statistics**

| | 50 | 70 | 90 |
|---|---|---|---|
| HSapiens/all | 0.284 | 0.382 | 0.666 |
| HSapiens/median | 0.139 | 0.274 | 0.630 |
| HSapiens/mean | 0.071 | 0.344 | 0.877 |

Table S3: **HSapiens: CDF pLDDT**

##### S1.1.2 PAeruginosa

As reported in Figure S3 and Table S4:

- Worst cases, top 3: ['File PAeruginosa/AF-Q9I728-F1-model_v4.pdb has 198 a.a.; pLDDT min/med/-max: 20.78 25.56 47.94', 'File PAeruginosa/AF-Q9I2K8-F1-model_v4.pdb has 327 a.a.; pLDDT min/med/-max: 20.37 26.57 89.72', 'File PAeruginosa/AF-Q9I665-F1-model_v4.pdb has 224 a.a.; pLDDT min/med/-max: 21.50 29.81 73.31']

- Median cases, 3 of them: ['File PAeruginosa/AF-Q9HTB7-F1-model_v4.pdb has 479 a.a.; pLDDT min/med/max: 47.61 95.54 98.73', 'File PAeruginosa/AF-Q9HVR7-F1-model_v4.pdb has 303 a.a.; pLDDT min/med/max: 39.46 95.54 98.50', 'File PAeruginosa/AF-Q9HXS6-F1-model_v4.pdb has 267 a.a.; pLDDT min/med/max: 37.32 95.54 98.81']

- Top cases, top 3: ['File PAeruginosa/AF-Q9HTP2-F1-model_v4.pdb has 497 a.a.; pLDDT min/med/-max: 54.80 98.85 98.98', 'File PAeruginosa/AF-Q9HWU0-F1-model_v4.pdb has 461 a.a.; pLDDT min/med/max: 42.05 98.85 98.98', 'File PAeruginosa/AF-Q9I121-F1-model_v4.pdb has 159 a.a.; pLDDT min/med/max: 90.82 98.86 98.96']

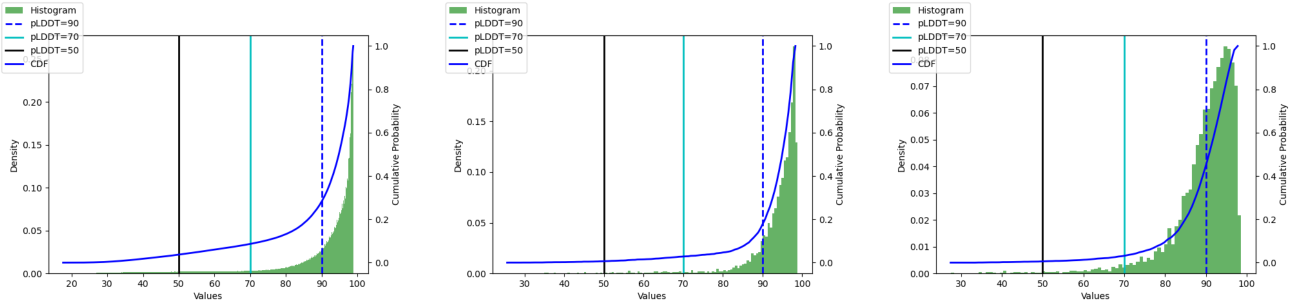

pLDDT all　　pLDDT median　　pLDDT mean

Figure S3: **PAeruginosa: pLDDT statistics**

| | 50 | 70 | 90 |
|---|---|---|---|
| PAeruginosa/all | 0.036 | 0.086 | 0.278 |
| PAeruginosa/median | 0.006 | 0.028 | 0.156 |
| PAeruginosa/mean | 0.005 | 0.030 | 0.396 |

Table S4: **PAeruginosa: CDF pLDDT**

##### S1.1.3 PFalciparum

As reported in Figure S4 and Table S5:

- Worst cases, top 3: ['File PFalciparum/AF-Q8ILT4-F1-model_v4.pdb has 2280 a.a.; pLDDT min/med/-max: 15.91 23.66 96.49', 'File PFalciparum/AF-C0H4Q1-F1-model_v4.pdb has 2580 a.a.; pLDDT min/med/max: 17.34 24.41 96.81', 'File PFalciparum/AF-C6KT74-F1-model_v4.pdb has 2404 a.a.; pLDDT min/med/max: 16.33 24.84 98.69']
- Median cases, 3 of them: ['File PFalciparum/AF-Q8IFL1-F1-model_v4.pdb has 231 a.a.; pLDDT min/med/max: 33.35 76.56 93.89', 'File PFalciparum/AF-Q8I419-F1-model_v4.pdb has 366 a.a.; pLDDT min/med/max: 27.79 76.59 97.05', 'File PFalciparum/AF-Q8IIM7-F1-model_v4.pdb has 698 a.a.; pLDDT min/med/max: 22.18 76.60 95.19']
- Top cases, top 3: ['File PFalciparum/AF-Q8IKK7-F1-model_v4.pdb has 337 a.a.; pLDDT min/med/-max: 69.22 98.66 98.94', 'File PFalciparum/AF-Q8IAY6-F1-model_v4.pdb has 198 a.a.; pLDDT min/med/max: 66.94 98.69 98.94', 'File PFalciparum/AF-Q8I3X4-F1-model_v4.pdb has 245 a.a.; pLDDT min/med/max: 49.70 98.76 98.97']

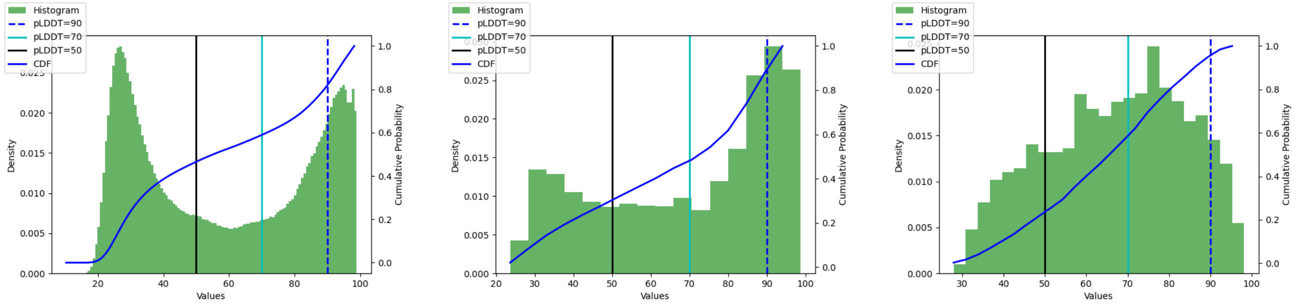

pLDDT all　　pLDDT median　　pLDDT mean

Figure S4: **PFalciparum: pLDDT statistics**

| | 50 | 70 | 90 |
|---|---|---|---|
| PFalciparum/all | 0.460 | 0.584 | 0.804 |
| PFalciparum/median | 0.261 | 0.441 | 0.757 |
| PFalciparum/mean | 0.197 | 0.529 | 0.915 |

Table S5: **PFalciparum: CDF pLDDT**
